# Microbial communities in retail draft beers and the biofilms they produce

**DOI:** 10.1101/2021.08.18.456920

**Authors:** Nikhil Bose, Daniel P. Auvil, Erica L. Moore, Sean D. Moore

## Abstract

In the beer brewing industry, microbial spoilage presents a consistent threat that must be monitored and controlled to ensure the palatability of a finished product. Many of the predominant beer spoilage microbes have been identified and characterized, but the mechanisms of contamination and persistence remain an open area of study. Post-production, many beers are distributed as kegs that are attached to draft delivery systems in retail settings where ample opportunities for microbial spoilage are present. As such, restaurants and bars can experience substantial costs and downtime for cleaning when beer draft lines become heavily contaminated. Spoilage monitoring on the retail side of the beer industry is often overlooked, yet this arena may represent one of the largest threats to the profitability of a beer if its flavor profile becomes substantially distorted. In this study, we sampled and cultured microbial communities found in beers dispensed from a retail draft system to identify the contaminating bacteria and yeasts. We also evaluated their capability to establish new biofilms in a controlled setting. Among four tested beer types, we identified over a hundred different contaminant bacteria and nearly twenty wild yeasts. The culturing experiments demonstrated that most of these microbes were viable and capable of joining new biofilm communities. From these data, we provide an important starting point for the efficient monitoring of beer spoilage in draft systems and provide suggestions for cleaning protocol improvements that can benefit the retail community.

**Importance:** Beer production, packaging, and service are each vulnerable to contamination by microbes that metabolize beer chemicals and impart undesirable flavors, which can result in the disposal of entire batches. Therefore, great effort is taken by brewmasters to reduce and monitor contamination during production and packaging. A commonly overlooked quality control stage of a beer supply chain is at the retail service end, where beer kegs supply draft lines in bars and restaurants under non-sterile conditions. We found that retail draft line contamination is rampant and that routine line cleaning methods are insufficient to efficiently suppress beer spoilage. Thus, many customers unknowingly experience spoiled versions of the beers they consume. This study identified the bacteria and yeast that were resident in draft beer samples and also assessed their abilities to colonize tubing material as members of stable biofilm communities.

## Introduction

Beer production involves controlled fermentation of plant sugar extracts in the presence of flavoring compounds to generate desirable beverages. During the brewing process, substantial effort is given to minimize exposure to spoilage microbes that compete for resources and impart undesirable flavors. In addition to careful fermentation, many finished beers are also filtered or pasteurized to further improve product stability, sometimes at the cost of product flavor quality. Unfortunately, these efforts go to waste if a beer is spoiled during the packaging, distribution, or dispensing stages. In our study, we observed that routine draft line cleaning procedures are insufficient to maintain beer quality in retail draft systems and that resilient microbial biofilms persist that rapidly reestablish complex spoilage communities. To begin to address this issue, we characterized microbial communities obtained from commercial draft beers and monitored their populations after they established biofilms during culturing.

In the nineteenth century, Louis Pasteur established that certain yeasts could be isolated and used to produce wines and beers with consistent and desirable characteristics (1). With those studies also came the discovery that beer and wine spoilage was caused by different microbes that competed for food resources and generated undesirable metabolites, such as lactic and acetic acid (1). Thus, the industry of fermented beverage production rapidly shifted away from so called “wild” inoculations and industry standards were put in place to carefully monitor and control the presence of both desirable and undesirable microbes (2). In the last few decades, the craft beer industry has revisited the use of alternative microbes and combinatorial culturing to greatly expand the style range and flavor profiles. With some irony, one goal of these efforts is to develop create products with scent and flavor complexities that match wild-fermented ales and lambics (3, 4). Nevertheless, great care and expense is still applied to minimize contamination by spoilage microbes and to ensure product stability (2, 5, 6).

Spoilage microbes enter the brewing process primarily from the addition of non-sterile ingredients, air exposure, or contaminated equipment. Several spoilage microbes are well known to the brewing community because they are commonly encountered and present a consistent threat; among these lactic acid bacteria (LAB), acetic acid bacteria (AAB), and wild yeasts represent dominant cohorts (6–10). Interestingly, these types of microbes are also present as desirable members of the microbial communities found in wild fermentations, wherein they can improve flavor balance and impart sour characteristics as the beers are aged to maturity (11–13). In these aging processes, groups of microbes overtake one another to dominate the community in a cascading fashion, with each group consuming old metabolites and creating new ones. In addition, members of a microbial community can exhibit synergistic or antagonistic relationships with each other, which promotes unpredictable community restructuring depending on the metabolic and combat capabilities of the founding members (14–18). The transitions through community structures are a key feature that provides unique complexity to the finished products. However, this type of conditioning process can be highly unpredictable; even different strains of a microbial species can exhibit notably different growth capabilities and differentially consume or release metabolites that alter beer flavor (4, 12).

Historically, the identification of spoilage microbes relied on the ability to culture a contaminant so that it could be subsequently characterized phenotypically and biochemically (7). More recently, sensitive techniques to detect known spoilage microbes have been developed that can reveal the presence and abundance of a particular microbe’s genome that employ either image cytometry (19), polymerase chain reactions (PCR) (8), bioluminescence (20), or molecular probing (21). While PCR is excellent for characterizing microbes in post-production beer and for predicting shelf life, it is unable to detect genes outside those that are targeted, so other potential spoilage microbes go unnoticed. These limitations could largely be overcome using next-generation DNA deep-sequencing to monitor mixed microbial communities because all recoverable DNA can be interrogated and the abundance of non-culturable microbes can also be established (22, 23). Unfortunately, the time and costs associated with deep-sequencing are not compatible with routine beer production protocols.

Deep-sequencing has been applied to thoroughly evaluate the presence of microbes and particular genes associated with spoilage in an active brewery (9). What emerged from that study were mosaic maps of microbial communities that were influenced both by location and nutrient availability in each brewery station. A main conclusion from that investigation was that repeated exposure to the beer itself was correlated with the abundance of genes that are associated with resistance to iso-alpha acids derived from hops. Thus, in addition to nutrient availability, the chemistry of a given beer, competition or predation by other microbes, and evasion from cleaning protocols become key aspects governing a beer spoilage microbial community.

In this study, we recovered the microbial communities from four different beer samples (starters) at two timepoints from a retail draft system and used DNA deep-sequencing to determine the relative abundances of the microbial genomes in each sample. In addition, each sample was used to inoculate experimental cultures with a nutrient-rich growth environment to invoke beer spoilage in the presence of draft line plastic plugs. The resulting microbial communities of the non-adherent (planktonic) fractions and the stably plastic-associated (biofilm) fractions were subsequently processed for deep-sequencing to establish the relative abundances of the microbes. All together, we detected 119 bacterial and 18 fungal species as contaminants in these draft systems. The samples collected at two different time points yielded different bacterial communities in the same beers. We also identified members of these starter communities that re-established themselves as members of new biofilms and we were able to identify bacteria that preferred growth in biofilms. Therefore, this study broadens the understanding of beer spoilage beyond controlled brewery settings and sets a foundation for improved retail service education and spoilage monitoring.

## Results

### Establishing a test platform for beer microbiota

Four retail beer draft taps were selected for study that delivered a <u>l</u>ager (L), an <u>I</u>ndia pale ale (I), a <u>h</u>efeweizen (H), and an <u>e</u>xtra pale ale (E) (**Fig. 1A**). Aliquots of each draft sample were used to inoculate three replicate cultures. The selected growth medium was the same brand of lager drawn from the tap, but sourced from a can to avoid prior microbial contamination. To ensure sterility, the growth beer was also filter sterilized prior to delivery into the culture tubes. To provide a growth surface that appropriately represented the beer service lines in this system, the experimental growth tubes also contained uniformly-dimensioned plugs of draft line tubing that rested at an angle within the culture medium to allow non-biofilm settling microbes to drift to the tube bottoms (**Fig. 1B**). The plug surface area and liquid volume was the same for each replicate. Upon sealing, no additional atmospheric exposures occurred until harvesting. After one week, the samples were mixed to re-distribute the microbes and they were incubated for an additional week.

**Figure 1.**
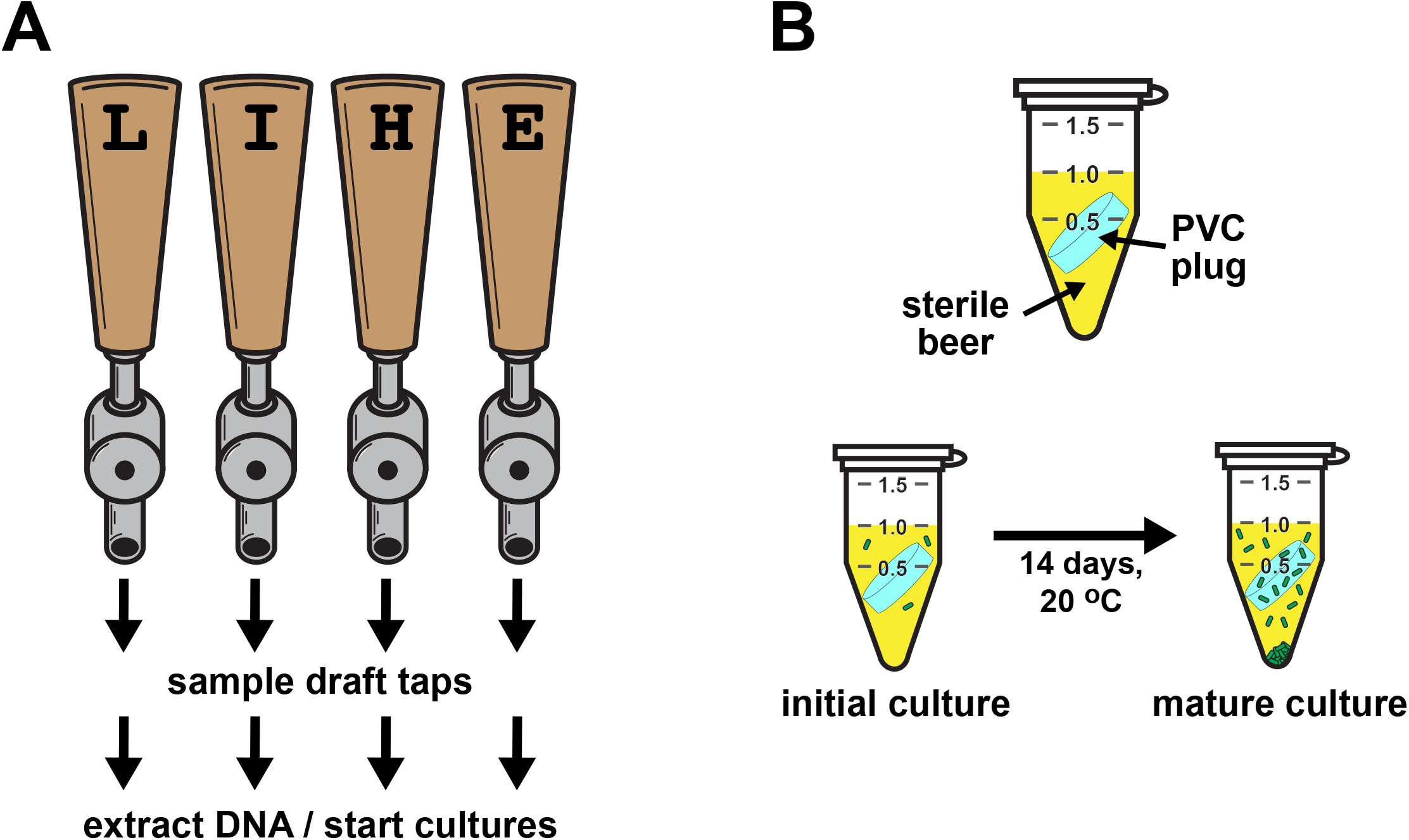
Beer sampling and biofilm development. A retail setting was chosen that served draft beers with diverse styles. **A)** Beer samples were collected from four draft taps serving a <u>l</u>ager (L), an <u>I</u>PA (I), a <u>h</u>efeweizen (H), or an <u>E</u>PA (E) as the first draws of that day. The microbes in each were used either for direct DNA extractions or for inoculating cultures. **B)** Each beer starter was used to inoculate three tubes containing sterile lager beer as a growth medium and uniform plugs of draft line plastic. The cultures were allowed to develop for two weeks before extracting DNA from the planktonic and biofilm cells. Approximately one year later, a second sampling was performed from the same taps (which were still serving the same beers) and the experiment was repeated.

### Bacterial diversity in the starter samples

The samples were processed to recover total DNA from the free living (planktonic) and adherent (biofilm) cells, and polymerase chain reactions (PCR) were used to amplify either the hypervariable V3-V4 regions of the bacterial 16S ribosomal genes or fungal ITS2 regions (24, 25). These PCR amplicons were then barcoded, pooled, and sequenced using paired-end Illumina technology (**Supporting Data S1**) (26). To evaluate bacterial communities, the 16S V3-V4 sequences were computationally processed to identify the source species and their relative abundances in the samples. In the ‘starter’ samples collected from the draft taps in the first year, each beer had a different and very rich community structure. In these beers, we identified 164 different V3-V4 sequences that were derived from 98 species of bacteria (**Data S2**). For the samples collected in the second year, we identified 143 unique sequences derived from 72 species, most of which were the same as those observed in the year 1 collection (**Data S3**).

Most bacteria contain multiple copies of their 16S genes and the V3-V4 regions within them may differ in a single organism (27, 28). Moreover, sub-species (strains) of bacteria frequently have the same V3-V4 regions as other members of the species (28, 29). Therefore, while the total counts of sequences and putative species names are informative for comparing community structures, they are not directly reflective of the number of different strains. Nonetheless, we gained an important insight into the diversity of bacteria that can be present in retail draft beers.

By comparing sample V3-V4 sequences to database 16S sequences, we were able to confidently assign identities to nearly all bacteria at the genus level and to the species level for a smaller cohort (**Data S2**). To reduce this large group for discussion purposes, we computationally filtered the collection of sequences for those that were also present either in each of the three biofilm or in each of the three planktonic culture replicates derived from them. This processing reduced the number of genera to 31 for the year 1 samples and 12 for the year 2 sample. These selected genera were then compared for their evolutionary relatedness within the eubacterial kingdom (**Fig. 2A**). During this analysis, we discovered that approximately half of these genera were recovered predominantly from either the biofilm or planktonic culture samples, suggesting that those bacteria had preferred biological niches in our growth experiments (**Fig. 2A**).

**Figure 2.**
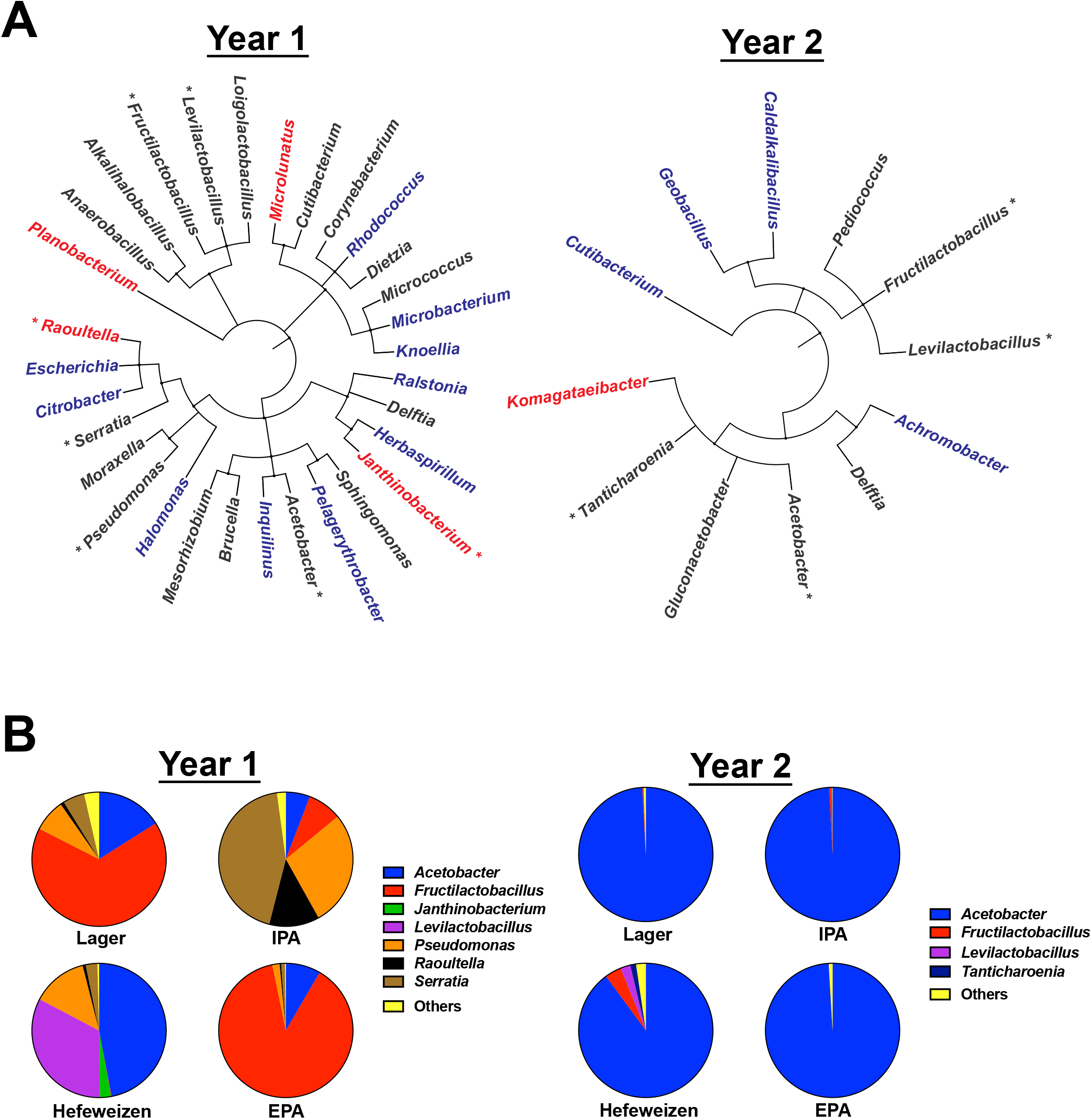
Bacteria present in the starter samples. The V3-V4 hypervariable regions of bacterial 16S rRNA genes from all project samples were amplified using PCR and pooled prior to Illumina paired-end sequencing. Zero-radius operational taxonomic units (zOTUs) were assigned to the sequence collection and the identities of the source bacteria were determined to the genus level. **A)** Taxonomic bushes illustrate the diversity of bacteria present both in a starter beer and in a cultured sample; plotted from kingdom to genus for each year. Sequences present at >1% are marked with asterisks. Cultured bacteria that were detected predominantly in biofilms are colored blue and those detected predominantly as planktonic are colored red. **B)** Pie-charts of sequence reads for genera present at >1% abundance in the initial samples (lesser abundances were grouped together as “others”).

To gain insight into the dominant community members in this group, we identified those bacteria whose sequence reads were present at greater than 1% of the total reads in any given starter sample (**Fig. 2B**). This comparison of the community structures revealed that the most dominant members in the year 1 collection varied substantially between each style of beer, with *Acetobacter*, *Fructilactobacillus*, or *Serratia* as major members. The observed abundances in the year 2 samples were markedly different, with each sample being dominated by *Acetobacter*. These comparisons highlight an important conclusion from this study: although major community members were similar between the first- and second-year collections, the relative abundances of the bacteria changed dramatically between sampling events. Thus, retail draft line communities are dynamic and there is no particular pattern of bacterial abundance that could predict which beer they came from.

### Preferences for biofilm or planktonic growth

The conclusion that some bacteria preferred occupancy in either the biofilm or planktonic communities was derived from a relatively straightforward visual inspection of the read count data for each growth experiment. However, that analysis overlooks bacteria that were abundant in both communities whose relative proportions in the biofilm and planktonic communities differed significantly after culturing. Bacteria that show a propensity to grow better in a biofilm relative to other community members may be capable of dominating when biofilms are re-established after draft line cleaning. To identify bacteria that exhibited such a behavior in our cultures, we applied an analytical method that compares the abundance of a given sequence read relative to a reference sequence that was present in all samples (30). This analysis has the advantage in that it does not require counting the absolute numbers of microbes in a given sample, which is intractable in biofilm studies. We elected to use the *Acetobacter* sequence (zOTU1) as a reference because it was abundant and present in all data sets. We were also able to leverage the outcomes of the replicated cultures to reveal reproducible behaviors.

For this analysis, we first filtered the data sets to only include sequence reads (zOTUs) that were present at greater that 0.1% compared to the reference sequence in each of the samples (**Data S2**). This filtering strategy avoided a pitfall caused by low abundance sequences: small changes in read counts between samples can be incorrectly perceived as very large changes in relative abundance. We then calculated how the relative abundance of a sequence read in the cultured biofilm or planktonic samples changed compared its abundance in the starter (relative differentials) (30). Taking a log of those values provides an easier interpretation of any changes because increases or decreases become evenly distributed around zero (which is no change). The relative differentials were averaged across the three replicates and comparisons were made between biofilm and planktonic residency. For example, of the 14 sequence sets (the starter, the three biofilm, and the three planktonic) that passed the abundance filter in the year 1 lager culture experiments, there were seven instances where the read abundances were significantly different between the biofilm and planktonic environments (**Fig. 3A and 3B**). In this representation, a positive value for the difference between biofilm abundance and planktonic abundance, the delta, means that the bacterium was more prevalent in that biofilm community with respect to the reference.

**Figure 3.**
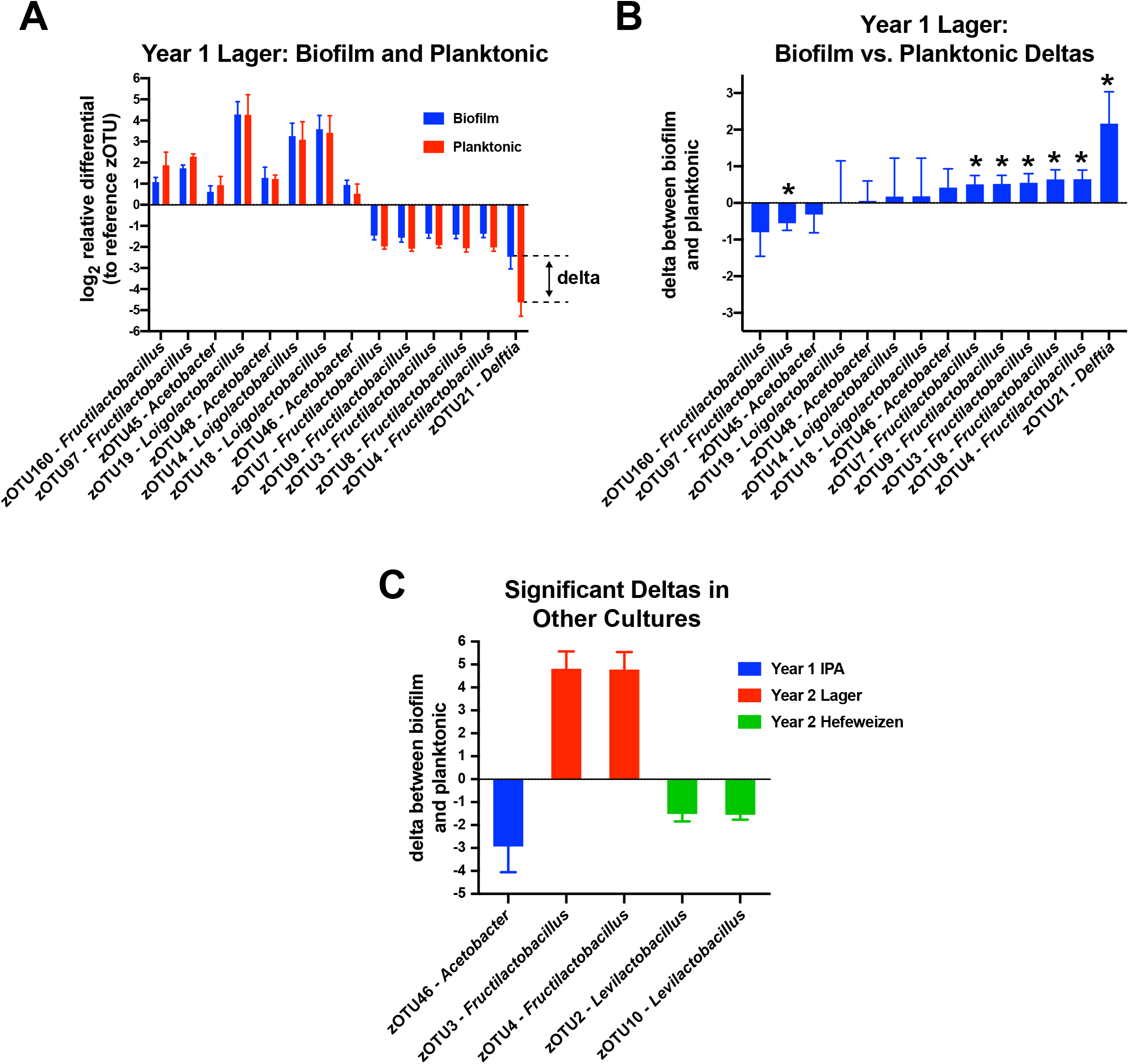
Evaluating bacterial biofilm preferences. The sequence read counts in each sample were used to calculate relative abundances with respect to a common reference sequence in each sample. Those ratios were then used to establish changes in their relative abundances in the biofilm or planktonic communities compared to their abundances in the starter samples. **A)** Bar plots of the log2 transforms of the relative differentials for bacteria in the lager cultures (a two-fold change is one unit on the ordinate axis). Error bars indicate the standard deviations between the three culture replicates. The ‘delta’ is the difference between the biofilm change and the planktonic change. **B)** Bar plots of the deltas, with negative values indicating a preference for the planktonic niche and positive values for the biofilms. Delta values from pairs that had significant differences between the biofilm and planktonic groups are marked with asterisks (*t*-test P values <0.05). **C)** Five additional significant biofilm deltas observed among the other seven beer experiments.

Strong additional support for this analytical approach came from an interesting discovery we made regarding the five dominant *Fructilactobacillus lindneri* sequences in the year 1 lager experiment (zOTUs 3,4,7,8, and 9), where each was significantly overrepresented in those biofilms by ~50% (**Fig. 3B**). *F. lindneri* has seven 16S genes and the read counts among the samples for those zOTUs had nearly consistent proportions in all samples of 2:2:1:1:1, respectively. This correlation suggests there was a dominant *F. lindneri* strain with two copies of a 16S gene containing the zOTU3 or zOTU4 sequence, and three 16S genes with each of the others. A similar phenomenon was observed with *Loigolactobacillus backii*, where its five 16S genes appeared in a 3:1:1 ratio (zOTUs 14, 18, and 19), indicating that three 16S sequences were identical and remaining two were different. An *L. backii* representative genome (NCBI RefSeq ID# NZ_CP014873.1) also shows a 3:1:1 ratio of V3-V4 region sequences. In contrast to *F. lindneri*, the log-ratio comparisons revealed that these *L. backii* sequences consistently had positive values, although with very similar deltas between the biofilm and planktonic fractions. These results indicate that the abundances of these 16S genes were changing in consistent proportions regardless of their copy number, which would be a requirement if they were members of the same genome. Moreover, this type of analysis may be useful in future studies for teasing out the number of strains within samples that have similar or shared gene sequences.

We performed the filtering and delta analysis for the seven other culturing experiments (three beers from year 1 and four from year 2) and identified five sequences that also exhibited significant deviations between the biofilm and planktonic samples (**Fig. 3C**) (**Data S2 and S3**). Interestingly, the *F. lindneri* zOTU3 and zOTU4 were also overrepresented in the year 2 lager biofilms (by ~30-fold) and the corresponding zOTUs 7, 8, and 9 were again present in the same ratios as was observed in the year 1 experiment. Unfortunately, the read counts of the latter three were too low in the starter sample to survive our filtering protocol. Nonetheless, this organism grew well in the lager and it became proportionally more abundant in the biofilms.

### Fungal diversity

In addition to bacteria, beers are known to be susceptible to contamination by so called ‘wild’ yeasts, which are non-conventional or uncultivated members of the fungi kingdom. Therefore, we sought to additionally search within our DNA extracts for sequences derived from fungi in an effort to characterize their roles in the contamination communities. PCR was used to amplify hypervariable segments of fungal genomes between the 5.8S and 28S rRNA genes (termed the internal transcribed spacer 2 region, ITS2) and then sequenced and computationally analyzed in a similar fashion to the analysis of the bacterial sequences (**Data S4** and **S5**).

We obtained PCR amplicons from each of the samples from the year 1 project stage, but were not able to recover amplicons from the year 2 samples, which suggests fungi were in very low abundance relative to the bacteria. Within the year 1 samples, we identified 18 ITS2 fungal sequences and were able to confidently assign 16 species as the sources of them. There is no method to correct for ITS2 copy numbers in fungal genomes, which can have tens or thousands of copies, and which can be variable within a given species or strain (31–33). Therefore, as with the bacterial study, the read counts should not be considered to directly correlate with cellular abundance.

The fungal species discovered in the starting beers are shown in **Fig. 4**. As with the bacterial communities among these samples, each beer exhibited a notably different fungal community distribution. A highly abundant *Saccaromyces cerevisiae* sequence (zOTU1) was present in the hefeweizen sample, which was anticipated because hefeweizens are not filtered prior to service and they are visibly turbid from the brewing yeast. Without additional sequence information on the genomes of these organisms, we are unable to determine if the prominent *S. cerevisiae* sequence is the same among the other samples because this ITS2 sequence is the same in many strains. The other species are considered to have been beer contaminants.

**Figure 4.**
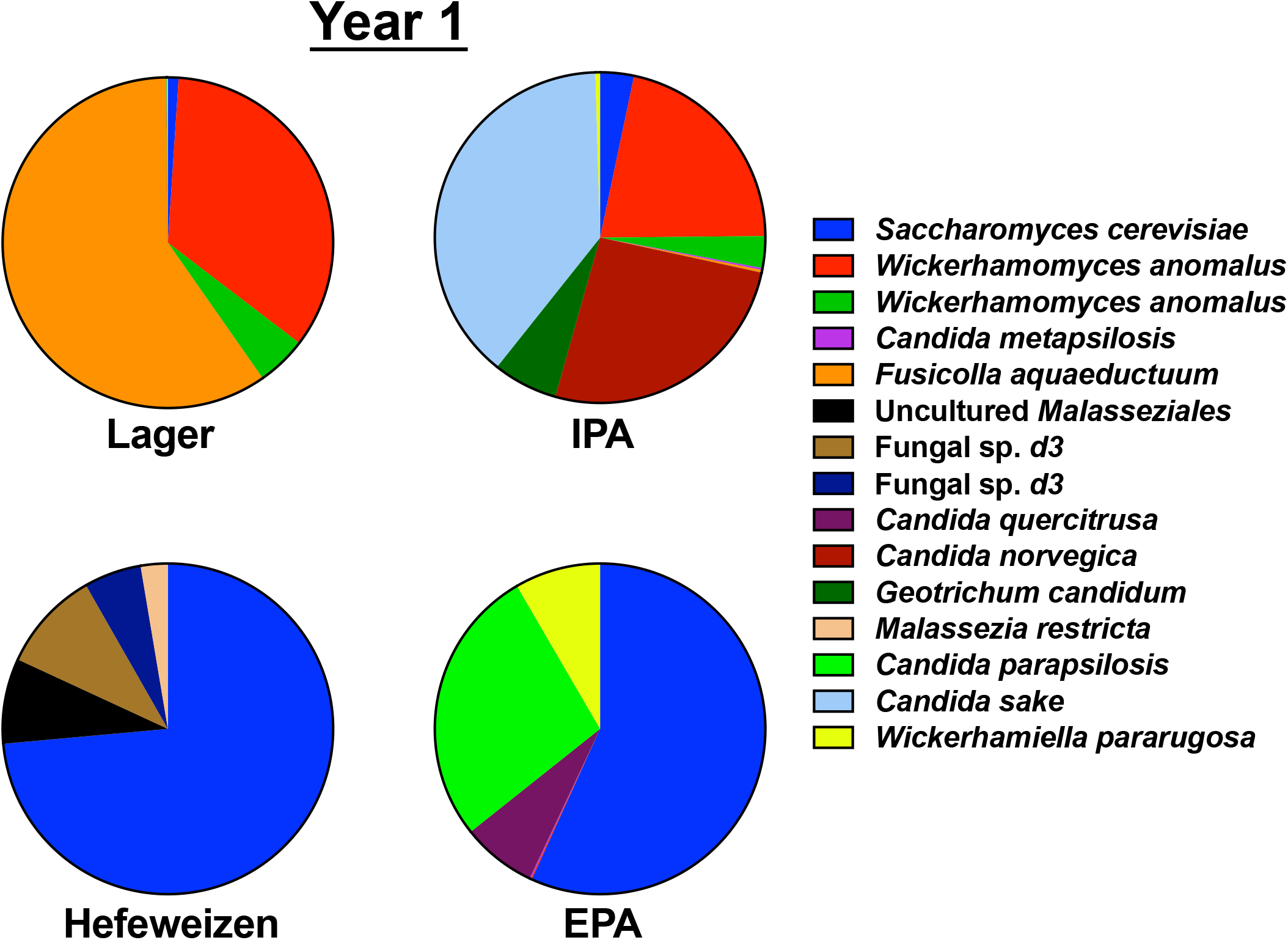
Fungi present in the samples. The ITS2 hypervariable regions between the fungal 5.8S and 28S rRNA genes were sequenced from the year 1 samples and processed to reveal abundance and to identify the organisms. Pie-charts show zOTU abundances for reads present at greater than 1% in each of the four sampled beers, with rarer reads grouped as ‘others’. The dominant *S. cerevisiae* sequence (zOTU1) that was abundant in the hefeweizen beer was likely from the brewing yeast, which is not removed from this style of beer; however, it was detected in all starter samples.

Only a few of the yeasts grew well in the subsequent culturing experiments and none of those outgrowths exhibited a significant bias between the biofilm and planktonic fractions. In the samples inoculated with the lager microbes, the *S. cerevisiae* and *Wickerhamomyces anomalus* strains grew well in all three cultures (**Data S5**). The *Fusicolla aquaeductuum* that provided the most sequence reads in this starter sample was not detected in five of the extracted samples and there were only a few reads in a planktonic sample that likely remained from the inoculum without any culture growth. In the cultures inoculated with the IPA starter, *W. anomalus* again grew well along with *Candida metapsilosis*, which was not abundant in this starter. The *Candida sake* that was dominant in this starter did not grow. Of the five prominent yeasts in the hefeweizen starter, only the *S. cerevisiae* grew out in the cultures. Interestingly, a *Brettanomyces* strain that was not detected in this starter was detected in all of the cultured samples. Its absence in the data for the starter was likely caused by the dominance of the *S. cerevisiae* sequence reads in that starter sample, but there was no correlation between the abundance of *S. cerevisiae* and *Brettanomyces* in the cultures. Finally, of the four dominant yeasts in the EPA starter, only the *S. cerevisiae* grew out in the cultures. These observations indicate that, unlike the recovered bacteria, there was a stark difference between the yeasts that were presence in the starter and their capability to grow in the cultures.

Overall, the bacterial and fungal community structures that were present in these four starters were not maintained in the lab cultures, even for the cultures grown using the same lager brand. Also, the communities in each draft line became restructured over the course of a year and the dominant members changed, which indicates that the chemistry of the beers themselves was not fully defining the community structures. These observations highlight an important value in ‘field monitoring’ microbial communities in different industrial or clinical settings because single snapshots of diversity in complex microbial communities neither reflect their history nor predict their futures.

## Discussion

We have provided a comprehensive survey of the microbial communities that inhabit retail draft beers and their ability to establish new communities in a controlled setting. In the cases of the two lager experiments, the microbes were presented with the same nutrient environment as their source environment, whereas the other communities were forced to adapt from their source environment into the lager. At first glance, it may seem surprising that the communities became so different from their starters when cultured. However, the starter communities themselves were recovered from environments that were already in flux: because they were recovered as the first draft pours of the day, they were in the process of consuming leftover resources in the beer lines and they had not been supplied fresh nutrients for ~12 hours or more, depending on the last time beer was pulled completely through each system. In addition, the lab culturing was done on a long timescale and the dynamics of the communities over that time are unknown. The choice of a long culturing stage was motivated by our experiences with isolated cultures of *Acetobacter* and *Lactobacillus*, which can take several days or more to reach saturation in established culture media or sterile beer. Nonetheless, we were able to evaluate outgrowth and to identify bacteria with preferences for biofilm and planktonic growth. We suspect the microbes with a penchant for biofilm residence are likely to be members of the microbial cohorts that help to re-establish communities after cleaning events.

It is not surprising that many of the predominant members of the retail draft line communities are well known for causing beer spoilage in breweries, such as LABs and AABs. LABs are considered to be one of the dominant contaminant microbes in breweries and they can contain genes that render them hop-resistant, so they can persist in beers with high hop extract content (10, 34, 35). In a comprehensive survey of brewery microbes found on ingredients, exposed surfaces, and in the liquids of different brewing stages (9), it was revealed that bacteria in the family *Lactobacillaceae* had a high relative abundance in beer samples, but much less occupancy elsewhere in the brewery. This disparity likely reflects the fact that these bacteria are usually anaerobes and grow poorly or not at all when exposed to the open atmosphere (36). The methodology of the brewery study limited the taxonomic assignments to the family level; however, in our analysis we could confidently assign a species to strains of *Fructilactobacillus lindneri*, *Levilactobacillus brevis*, *Loigolactobacillus backii*, and *Pediococcus damnosus*. In another study, Although these microbes routinely spoil beers and are frequently consumed, they are not considered to be human pathogens (37, 38).

A recent investigation of the growth properties of *F. lindneri* indicated that this bacterium can enter a “viable but nonculturable” (VBNC) state at low temperatures and that it requires anaerobic conditions for robust growth (39). Therefore, it is missed by routine colony screening for contamination and can persist undetected in refrigerated beers for long periods of time. That study also revealed that *F. lindneri* cells can produce high levels lactic acid, acetic acid, and diacetyl as waste metabolites, even from the VBNC state. The detection of this bacterium in all of our beer samples and the robust growth the lager cultures indicates that it is likely a major contributor to the off-flavors encountered in the starter beer collection. Our finding that it also prefers to reside in the biofilm niche suggests it may be one of the community members that evades cleaning protocols. A convenient strategy to monitor the presence of this microbe in the retail setting would be useful for quality control.

The *Levilactobacillus brevis* strains identified in our study also belong to another known beer spoilage group (40, 41). They were abundant in the hefeweizen samples, but at least one was detected in each starter sample. Some strains of this bacterium have been well characterized because they are considered probiotic and are used in the production of yogurts, kimchi, sourdoughs, and other fermented foods (42–46). In addition to producing lactic acid, these bacteria tend to secrete copious amounts of exopolysaccharides that cause slimy textures and increased viscosity (47, 48). Thus, not only does their presence sour beers, but abundant growth causes unpleasant mouthfeel.

The *Acetobacter* strain that was the most prominent contaminant in our samples did not receive a species assignment for this study because although its V3-V4 sequence matches *A. farinalis*, it is also very similar to *A. orleanensis*, *A. persici*, and *A. cerevisiae*. Most of the characterized members of these species have been recovered from spoiled beers (49), and *A. farinalis* from fermented rice flour (50). As the genus name suggests, these bacteria produce acetic acid form a variety of sugars and ethanol, but they can also consume ethanol, acetic acid, and lactic acid as energy sources if oxygen is present (51, 52). The dominance of *Acetobacter* in each of the year 2 samples suggests there may have been an alteration in the maintenance of the draft lines that allowed more oxygen exposure. Limiting oxygen exposure is a well-established beer brewing mandate and we suggest consideration of this approach during draft line servicing.

Although the presented data were derived from cultures obtained at a single retail location, we have not identified any regional draft service system that does not contain substantial microbial contamination. So, what is a retailer to do? One obvious approach is to increase or change line cleaning events. However, we noted that when beer draft lines are cleaned, a common methodology is to remove the keg coupler, connect the line to a keg with cleaning solution, flush the line, and then reconnect the coupler back onto the same keg. If a beer in a keg or its valve was already contaminated, no amount of line cleaning will prevent regrowth of the community. Also, if multiple lines are cleaned using the same flushing keg, cross contamination between lines is very likely. Perhaps the couplers and keg valves could be disinfected during changes and cleaning. On top of this, there is no way to prevent contamination at the tap end: it is fully exposed to the far-from-aseptic bar environment.

Another consideration is the draft line material. Replacing lines is recommended from ‘time-to-time’, but there is no formal guideline on when, what material to use, or how to diagnose a serious biofilm problem. Cost and downtime are other major considerations. From this study, we can suggest that perhaps the draft beers be monitored for *Acetobacter* because they were common in all of the samples. We have recently isolated strains of *Acetobacter* from this draft system and are evaluating their ability to form biofilms on various materials. What we have so-far discovered is that they behave differently and produce substantially more biofilm than a reference *A. cerevisiae* strain we obtained from a stock center. That reference strain was recovered many decades ago from a brewery in Germany, long before modern draft line plastics were available. We suspect that the continued selection for bacteria and wild yeast to survive on a given surface during cleaning episodes has created collections of microbes that are tailored to be resilient in each environment. This is a completely analogous situation to the problems caused microbial biofilms in medical and industrial settings. For these reasons, additional biofilm growth and cleaning research is warranted. As indicated earlier, taking advantage of the 16S gene copy numbers in different bacteria may be useful in teasing out whether or not there are multiple strains and for validating conclusions drawn from the sequence read abundances.

This project was inspired by anecdotal observations of dramatic taste and odor changes in draft beers served at several local restaurants, a phenomenon we affectionately refer to as a ‘bowling alley’ taste. After preliminary plating and Sanger sequencing experiments, we recognized that the diversity of microbes was far greater than our expectation of a few dominant contaminants. This project was intended not only to expand our understanding of microbial community behaviors, but also to provide an impetus for brewers to consider the downstream influences on the perception of their beer quality: a customer trying a new beer will be off-put by spoilage metabolites and probably never realize that the flavor is substantially distorted. Likewise, bartenders are rarely trained (or allowed) to taste service beers, so they may never realize there is a problem with the product. Another aspect of contaminated beer is the potential for health impact. It is known that humans obtain their microbial flora from environmental exposure and that gut microbiology is dictated by diet (53). Therefore, this study provides a motivation for a formal study to be conducted to establish relationships between beer spoilage microbes, resident gut flora, and human physiology.

## Materials and Methods

### Sample collection and culturing

Approximately 200 mL of draft samples of four beers were collected into sterile cups: an American lager, an India Pale Ale, a hefeweizen, and an extra pale. After mixing by swirling, 50 mL was transferred to sterile conical tubes and placed on ice. Approximately 45 min later, each tube was centrifuged for 30 min at 3,500 RCF at 4 °C and 45 mL of the cleared supernatants was removed by aspiration. The pellets were resuspended in the residual 5 mL, creating 10X “starter” stocks. 1 mL aliquots of each sample were then prepared: one was frozen at −80 °C and the others were kept on ice to serve as inoculation sources that day. This process was repeated the following year from the same taps.

Solid substrates for biofilm development were formed by hand-stamp punching 0.25 inch plugs from 3/8 inch flexible polyvinylchloride tubing (Vinyl-Flex NFS-61, Advanced Technology Products, Milford Center, Ohio) and collected into a clean glass beaker. In a sterile laminar flow hood, the plugs were then submerged in a peracetic acid sterilizing solution (SporGon, Decon Labs Inc., King of Prussia, PA) for three hours, and then rinsed four times with 0.22 μm filter-sterilized HPLC-grade H2O. After the last wash, the residual water was drained, and each plug was transferred using sterile tweezers into sterile 1.7 mL microfuge tubes and stored until use.

Sterile growth medium was prepared by 0.22 μm filtering a commercial canned lager that was the same as the sampled American lager. 1 mL aliquots were then aseptically transferred to the tubes containing the PVC pellets. A set of 10 growth tubes was set aside as sterility controls and the remaining tubes were inoculated with 10 μL of the 10X microbe starter stock and mixed by vortexing. The culture tubes were then placed in a 25 °C incubator for two weeks with an additional mixing event after the first week.

### DNA extraction

Culture tubes were vortexed briefly to resuspend settled cells and a 500 μL aliquot was set aside as the planktonic cell fractions. The PVC plugs were then transferred with sterile tweezers to a microfuge tube containing 1 mL of 0.22 μm filter-sterilized HPLC-grade H_2_O and vortexed to remove non-adherent cells. This washing step was repeated two more times.

Bead ablation tubes were prepared by adding 100 μL of 0.1 mm zirconia beads (Research Products International, Mount Prospect, IL) to sterile 2 mL screw cap microcentrifuge tubes. 500 μL of a denaturing “extraction buffer” (5.5 M guanidinium thiocyanate, 100 mM potassium acetate, pH 5.5) was added to the beads before adding 200 μL of either planktonic cells or 200 μL of sterile water and a PVC plug. The tubes were then agitated twice using a FastPrep-24 lysis 5G instrument (MP Biomedicals, Irvine, CA) using the “Escherischia coli cells” setting. After disruption, cell debris and beads were collected by centrifugation at 14,000 RCF for 5 min and 500 μL of each cleared supernatant was transferred to a clean tube. 200 μL of isopropanol was added and mixed and the solution transferred to a DNA binding silica spin column (EconoSpin, Epoch Life Science, Missouri City, TX). After passing the solution through the column twice, the column was washed with 300 μL of extraction buffer, followed by three washes with “column wash buffer” (80 % ethanol, 10 mM Tris-Cl, 0.1 mM EDTA, pH 8.0). The columns were then dried by centrifugation and the samples eluted in 50 μL of “DNA buffer” (5 mM Tris-Cl, 0.1 mM EDTA, pH 8.0). DNA samples were stored at −20 °C.

### Library preparation and sequencing

Bacterial 16S V3-V4 regions were amplified by PCR using universal primers derived from DBact-0341-b-S-17 and S-D-Bact-0785-a-A-21 (24) containing unique adapters for the Nextera XT indexing kit according to the manufacturer’s instructions (Illumina Inc., San Diego, CA). Fungal ITS2 regions were amplified using separate unique adapter primers derived from IST3_KYO1 and IST4_KYO1 (25). After adding unique indices, each sample’s DNA concentration was determined using the Quant-iT PicoGreen kit (Molecular Probes, Inc., Eugene, OR) and equal mass portions of each were pooled prior to sequencing.

The amplicon pools were paired-end sequenced (250 cycles) using the MiSeq platform (Illumina Inc.) at either the Interdisciplinary Center for Biotechnology Research (year 1 data set, University of Florida, Gainesville, FL) or the UCF Genomics & Bioinformatics Cluster (year 2 data set, University of Central Florida, Orlando, FL).

### Sequence processing and bioinformatics

Trimmomatic was used to remove primer sequences and to filter out reads less than 150 bases long as well as reads with Phred quality (Q) scores less than 25 using a sliding window of 4 bases (54). VSEARCH was used to merge the forward and reverse sequences to generate the complete V3-V4 regions with a minimum total merged sequence length of 200 bases, minimum overlap length of 50 bases, and a maximum of 5 allowed mismatches across the alignment (55). Following merging, sequences with an expected error rate >1 were discarded. The remaining sequences were then dereplicated while counting the number of each unique sequence.

VSEARCH was used to denoise reads, remove chimeras, and to generate <u>z</u>ero-<u>r</u>adius <u>o</u>perational <u>t</u>axonomic <u>u</u>nits (zOTUs). Parameters for zOTU clustering included occurrence of a minimum of 50 unique reads within a 99% identity threshold. All error-filtered merged sequences were then mapped to zOTU sequences based on 99% alignment to generate a table containing sequence read counts per zOTU for each sample.

Taxonomic classification of zOTU sequences was carried out using the Bayesian Lowest Common Ancestry (BLCA) software, which employs full-length query-hit alignment scores to generate a weighted probability for taxonomic allocation (56). For bacterial classification, the NCBI 16S ribosomal RNA database was used (updated 06/24/2021) and for fungal classification the UNITE v6 database was configured for usage with BLCA (57). Classification was declared based on the lowest common ancestor with a cumulative posterior probability ≥ 80%. Fungal zOTUs were additionally manually compared to the NCBI’s non-redundant eukaryotic database using BLAST (58, 59).

The percent abundance of each zOTU was determined in each sample’s data set and used to calculate ratios relative to a common reference (zOTU1, which was present in all bacterial samples) as previously described (30). Finally, the log2 transforms of these reference frames were used to evaluate relative changes in the bacterial communities during culturing and to compare the relative abundances in biofilms and planktonic samples.

### Data analysis and graphics

Taxonomy bushes were generated using the Interactive Tree of Life (iTOL) (60). Data were sorted and processed using Excel (Microsoft, Redmond, WA). Data were plotted using Prism (Graphpad, San Diego, CA) and figures were generated using Illustrator (Adobe Inc., San Jose, CA).

## Acknowledgements

This work was partially supported by NIH grant 1R01GM118896 and a UCF ‘What’s Next’ Quality Enhancement Plan grant. The authors thank Anna Ward and Laurie Agosto for their technical assistance, as well as the following UCF undergraduates who participated in this project as members of the UCF Applied Industrial Microbiology program: Cesar Ballesteros, Taylor Croteau, Mahammad Gardashli, Haley Hardin, Christopher Hawkins, Sarah Kampiyil, Ashley Lima-Acosta, Jordan Palmer, Vanessa Parra-Gonzalez, Tristan Tran, Michael Tucker, Paola Andrade, Erick Bardalez, Danielle Beetler, Amy Freiberg, Chanelle Hunter, Hayden Kim, Imani Pascoe, Dana Ramsey, Keith Taylor, Evie Vincent, and Emily Wilson.

## Supporting Information Legends

**Supporting Data S1.** The bacterial V3-V4 zero-radius operational taxonomic unit (zOTU) nucleotide sequences in FASTA format.

**Supporting Data S2.** Bacterial V3-V4 read abundances in the year 1 starter samples. Illumina paired-end sequence reads of the bacterial V3-V4 amplicons were computationally processed to reveal abundances and taxons. The workbook contains several tab-separated sheets: “Year1 starter counts” contains the list of unique sequences (zOTUs) that were used to determine the number of species. Note that although many of the species assignments have low confidence, the putative names serve as placeholders that were used for counting. “Genera >1% Year1” lists the genera present in each starter beer if they were present in any of them at >1% of the total sequence reads. “Lager” contains the zOTUs designations, a sortable sequence ID, sortable taxonomies at different levels followed by the confidence scores assigned by the BLCA software, and the overall abundance in the entire project set. The original order of ascending zOTU numbering can be obtained by sorting the numerical “ID” column in ascending order. These columns are followed by an analysis of relative abundance in each planktonic sample, each biofilm sample, and the starter sample. The “Lager_Starter_Genus” and “Lager_Starter_Species” sheets contain the calculations for the summaries of abundances. The “Lager Abundance Filter” contains the calculations for establishing an abundance threshold and whether to include or exclude that zOTU from the subsequent biofilm/planktonic delta tests.

“Lager delta T-tests” contains the calculations of the averages of the log2 relative differentials and whether or not there is a statistically significant difference between the biofilm and planktonic community values (*p* </= 0.05). The additional tabs contain the same data analysis for each of the other three beers.

**Data S3.** Bacterial V3-V4 read abundances in the year 2 samples. This workbook has the same layout as the year 1 workbook.

**Data S4.** The fungal ITS2 zero-radius operational taxonomic unit (zOTU) sequences in FASTA format.

**Data S5.** Fungal ITS2 read abundances in the year 1 samples. Delta T-tests were only performed on samples with reads that passed the abundance filters.

## References

1. Barnett JA. 2000. A history of research on yeasts 2: Louis Pasteur and his contemporaries, 1850-1880. Yeast 16:755–71.

2. Bamforth CW. 2017. Progress in Brewing Science and Beer Production. Annu Rev Chem Biomol Eng 8:161–176.

3. Larroque MN, Carrau F, Fariña L, Boido E, Dellacassa E, Medina K. 2021. Effect of Saccharomyces and non-Saccharomyces native yeasts on beer aroma compounds. Int J Food Microbiol 337:108953.

4. Dysvik A, La Rosa SL, De Rouck G, Rukke EO, Westereng B, Wicklund T. 2020. Microbial Dynamics in Traditional and Modern Sour Beer Production. Appl Environ Microbiol 86.

5. Thesseling FA, Bircham PW, Mertens S, Voordeckers K, Verstrepen KJ. 2019. A Hands-On Guide to Brewing and Analyzing Beer in the Laboratory. Curr Protoc Microbiol 54:e91.

6. Wagner EM, Thalguter S, Wagner M, Rychli K. 2021. Presence of Microbial Contamination and Biofilms at a Beer Can Filling Production Line. J Food Prot 84:896–902.

7. Suzuki K. 2011. 125th Anniversary Review: Microbiological Instability of Beer Caused by Spoilage Bacteria. The Institute of Brewing & Distilling:131–155.

8. Janagama HK, Mai T, Han S, Nadala L, Nadala C, Samadpour M. 2018. Dipstick Assay for Rapid Detection of Beer Spoilage Organisms. J AOAC Int 101:1913–1919.

9. Bokulich NA, Bergsveinson J, Ziola B, Mills DA. 2015. Mapping microbial ecosystems and spoilage-gene flow in breweries highlights patterns of contamination and resistance. Elife 4.

10. Rodríguez-Saavedra M, González de Llano D, Moreno-Arribas MV. 2020. Beer spoilage lactic acid bacteria from craft brewery microbiota: Microbiological quality and food safety. Food Res Int 138:109762.

11. De Roos J, Verce M, Aerts M, Vandamme P, De Vuyst L. 2018. Temporal and Spatial Distribution of the Acetic Acid Bacterium Communities throughout the Wooden Casks Used for the Fermentation and Maturation of Lambic Beer Underlines Their Functional Role. Appl Environ Microbiol 84.

12. Spitaels F, Wieme AD, Janssens M, Aerts M, Daniel HM, Van Landschoot A, De Vuyst L, Vandamme P. 2014. The microbial diversity of traditional spontaneously fermented lambic beer. PLoS One 9:e95384.

13. Spitaels F, Wieme AD, Janssens M, Aerts M, Van Landschoot A, De Vuyst L, Vandamme P. 2015. The microbial diversity of an industrially produced lambic beer shares members of a traditionally produced one and reveals a core microbiota for lambic beer fermentation. Food Microbiol 49:23–32.

14. Basler M, Ho BT, Mekalanos JJ. 2013. Tit-for-tat: type VI secretion system counterattack during bacterial cell-cell interactions. Cell 152:884–94.

15. Chatterjee P, Sass G, Swietnicki W, Stevens DA. 2020. Review of Potential Pseudomonas Weaponry, Relevant to the Pseudomonas-Aspergillus Interplay, for the Mycology Community. J Fungi (Basel) 6.

16. Smith WPJ, Vettiger A, Winter J, Ryser T, Comstock LE, Basler M, Foster KR. 2020. The evolution of the type VI secretion system as a disintegration weapon. PLoS Biol 18:e3000720.

17. Thiery S, Kaimer C. 2020. The Predation Strategy of Myxococcus xanthus. Front Microbiol 11:2.

18. Torres-Guardado R, Esteve-Zarzoso B, Reguant C, Bordons A. 2021. Microbial interactions in alcoholic beverages. Int Microbiol doi:10.1007/s10123-021-00200-1.

19. Hodgkin M, Purseglove SM, Chan LL, Perry J, Bolton J. 2020. A novel image cytometry-based Lactobacillus bacterial enumeration method for the production of kettle sour beer. J Microbiol Methods 177:106031.

20. Takahashi T, Nakakita Y, Nakamura T. 2019. Rapid Single Cell Detection of Lactic Acid Bacteria in the Beer Using Bioluminescence Method. Biocontrol Sci 24:29–37.

21. Paradh AD, Hill AE, Mitchell WJ. 2014. Detection of beer spoilage bacteria Pectinatus and Megasphaera with acridinium ester labelled DNA probes using a hybridisation protection assay. J Microbiol Methods 96:25–34.

22. De Filippis F, Parente E, Ercolini D. 2017. Metagenomics insights into food fermentations. Microb Biotechnol 10:91–102.

23. Cao Y, Fanning S, Proos S, Jordan K, Srikumar S. 2017. A Review on the Applications of Next Generation Sequencing Technologies as Applied to Food-Related Microbiome Studies. Front Microbiol 8:1829.

24. Klindworth A, Pruesse E, Schweer T, Peplies J, Quast C, Horn M, Glöckner FO. 2013. Evaluation of general 16S ribosomal RNA gene PCR primers for classical and next-generation sequencing-based diversity studies. Nucleic Acids Res 41:e1.

25. Toju H, Tanabe AS, Yamamoto S, Sato H. 2012. High-coverage ITS primers for the DNA-based identification of ascomycetes and basidiomycetes in environmental samples. PLoS One 7:e40863.

26. Caporaso JG, Lauber CL, Walters WA, Berg-Lyons D, Huntley J, Fierer N, Owens SM, Betley J, Fraser L, Bauer M, Gormley N, Gilbert JA, Smith G, Knight R. 2012. Ultra-high-throughput microbial community analysis on the Illumina HiSeq and MiSeq platforms. ISME J 6:1621–4.

27. Acinas SG, Marcelino LA, Klepac-Ceraj V, Polz MF. 2004. Divergence and redundancy of 16S rRNA sequences in genomes with multiple rrn operons. J Bacteriol 186:2629–35.

28. Johnson JS, Spakowicz DJ, Hong BY, Petersen LM, Demkowicz P, Chen L, Leopold SR, Hanson BM, Agresta HO, Gerstein M, Sodergren E, Weinstock GM. 2019. Evaluation of 16S rRNA gene sequencing for species and strain-level microbiome analysis. Nat Commun 10:5029.

29. Lukjancenko O, Wassenaar TM, Ussery DW. 2010. Comparison of 61 sequenced Escherichia coli genomes. Microb Ecol 60:708–20.

30. Morton JT, Marotz C, Washburne A, Silverman J, Zaramela LS, Edlund A, Zengler K, Knight R. 2019. Establishing microbial composition measurement standards with reference frames. Nat Commun 10:2719.

31. Lofgren LA, Uehling JK, Branco S, Bruns TD, Martin F, Kennedy PG. 2019. Genome-based estimates of fungal rDNA copy number variation across phylogenetic scales and ecological lifestyles. Mol Ecol 28:721–730.

32. Peter J, De Chiara M, Friedrich A, Yue JX, Pflieger D, Bergström A, Sigwalt A, Barre B, Freel K, Llored A, Cruaud C, Labadie K, Aury JM, Istace B, Lebrigand K, Barbry P, Engelen S, Lemainque A, Wincker P, Liti G, Schacherer J. 2018. Genome evolution across 1,011 Saccharomyces cerevisiae isolates. Nature 556:339–344.

33. Gorter de Vries AR, Pronk JT, Daran JG. 2017. Industrial Relevance of Chromosomal Copy Number Variation in Saccharomyces Yeasts. Appl Environ Microbiol 83.

34. Feyereisen M, Mahony J, O’Sullivan T, Boer V, van Sinderen D. 2020. A Plasmid-Encoded Putative Glycosyltransferase Is Involved in Hop Tolerance and Beer Spoilage in Lactobacillus brevis. Appl Environ Microbiol 86.

35. Maifreni M, Frigo F, Bartolomeoli I, Buiatti S, Picon S, Marino M. 2015. Bacterial biofilm as a possible source of contamination in the microbrewery environment. Food Control 50:809–805.

36. Zheng J, Wittouck S, Salvetti E, Franz C, Harris HMB, Mattarelli P, O’Toole PW, Pot B, Vandamme P, Walter J, Watanabe K, Wuyts S, Felis GE, Gänzle MG, Lebeer S. 2020. A taxonomic note on the genus Lactobacillus: Description of 23 novel genera, emended description of the genus Lactobacillus Beijerinck 1901, and union of Lactobacillaceae and Leuconostocaceae. Int J Syst Evol Microbiol 70:2782–2858.

37. Ricci A, Allende A, Bolton D, Chemaly M, Davies R, Girones R, Herman L, Koutsoumanis K, Lindqvist R, Nørrung B, Robertson L, Ru G, Sanaa M, Simmons M, Skandamis P, Snary E, Speybroeck N, Ter Kuile B, Threlfall J, Wahlström H, Cocconcelli PS, Klein G, Prieto Maradona M, Querol A, Peixe L, Suarez JE, Sundh I, Vlak JM, Aguilera-Gómez M, Barizzone F, Brozzi R, Correia S, Heng L, Istace F, Lythgo C, Fernández Escámez PS. 2017. Scientific Opinion on the update of the list of QPS-recommended biological agents intentionally added to food or feed as notified to EFSA. Efsa j 15:e04664.

38. Lavermicocca P, Reguant C, Bautista-Gallego J. 2021. Editorial: Lactic Acid Bacteria Within the Food Industry: What Is New on Their Technological and Functional Role. Front Microbiol 12:711013.

39. Liu J, Li L, Li B, Peters BM, Deng Y, Xu Z, Shirtliff ME. 2017. First study on the formation and resuscitation of viable but nonculturable state and beer spoilage capability of Lactobacillus lindneri. Microb Pathog 107:219–224.

40. Jespersen L, Jakobsen M. 1996. Specific spoilage organisms in breweries and laboratory media for their detection. Int J Food Microbiol 33:139–55.

41. Munford ARG, Chaves RD, Granato D, Sant’Ana AS. 2020. Modeling the inactivation of Lactobacillus brevis DSM 6235 and retaining the viability of brewing pitching yeast submitted to acid and chlorine washing. Appl Microbiol Biotechnol 104:4071–4080.

42. Xiao T, Yan A, Huang JD, Jorgensen EM, Shah NP. 2020. Comparative Peptidomic and Metatranscriptomic Analyses Reveal Improved Gamma-Amino Butyric Acid Production Machinery in Levilactobacillus brevis Strain NPS-QW 145 Cocultured with Streptococcus thermophilus Strain ASCC1275 during Milk Fermentation. Appl Environ Microbiol 87.

43. Park E, Kim KT, Choi M, Lee Y, Paik HD. 2021. In Vivo Evaluation of Immune-Enhancing Activity of Red Gamju Fermented by Probiotic Levilactobacillus brevis KU15154 in Mice. Foods 10.

44. Somashekaraiah R, Mottawea W, Gunduraj A, Joshi U, Hammami R, Sreenivasa MY. 2021. Probiotic and Antifungal Attributes of Levilactobacillus brevis MYSN105, Isolated From an Indian Traditional Fermented Food Pozha. Front Microbiol 12:696267.

45. Bockwoldt JA, Fellermeier J, Steffens E, Vogel RF, Ehrmann MA. 2021. β-Glucan Production by Levilactobacillus brevis and Pediococcus claussenii for In Situ Enriched Rye and Wheat Sourdough Breads. Foods 10.

46. Abouloifa H, Khodaei N, Rokni Y, Karboune S, Brasca M, D’Hallewin G, Salah RB, Saalaoui E, Asehraou A. 2020. The prebiotics (Fructo-oligosaccharides and Xylo-oligosaccharides) modulate the probiotic properties of Lactiplantibacillus and Levilactobacillus strains isolated from traditional fermented olive. World J Microbiol Biotechnol 36:185.

47. Fraunhofer ME, Geissler AJ, Jakob F, Vogel RF. 2017. Multiple Genome Sequences of Exopolysaccharide-Producing, Brewery-Associated Lactobacillus brevis Strains. Genome Announc 5.

48. Fukao M, Zendo T, Inoue T, Fuke N, Moriuchi T, Yamane Y, Nakayama J, Sonomoto K, Fukaya T. 2019. Relation between cell-bound exopolysaccharide production via plasmid-encoded genes and rugose colony morphology in the probiotic Lactobacillus brevis KB290. Anim Sci J 90:1575–1580.

49. Cleenwerck I, Vandemeulebroecke K, Janssens D, Swings J. 2002. Re-examination of the genus Acetobacter, with descriptions of Acetobacter cerevisiae sp. nov. and Acetobacter malorum sp. nov. Int J Syst Evol Microbiol 52:1551–1558.

50. Tanasupawat S, Kommanee J, Yukphan P, Muramatsu Y, Nakagawa Y, Yamada Y. 2011. Acetobacter farinalis sp. nov., an acetic acid bacterium in the α-Proteobacteria. J Gen Appl Microbiol 57:159–67.

51. Cleenwerck I, De Vos P. 2008. Polyphasic taxonomy of acetic acid bacteria: an overview of the currently applied methodology. Int J Food Microbiol 125:2–14.

52. Gomes RJ, Borges MF, Rosa MF, Castro-Gómez RJH, Spinosa WA. 2018. Acetic Acid Bacteria in the Food Industry: Systematics, Characteristics and Applications. Food Technol Biotechnol 56:139–151.

53. David LA, Maurice CF, Carmody RN, Gootenberg DB, Button JE, Wolfe BE, Ling AV, Devlin AS, Varma Y, Fischbach MA, Biddinger SB, Dutton RJ, Turnbaugh PJ. 2014. Diet rapidly and reproducibly alters the human gut microbiome. Nature 505:559–63.

54. Bolger AM, Lohse M, Usadel B. 2014. Trimmomatic: a flexible trimmer for Illumina sequence data. Bioinformatics 30:2114–20.

55. Rognes T, Flouri T, Nichols B, Quince C, Mahé F. 2016. VSEARCH: a versatile open source tool for metagenomics. PeerJ 4:e2584.

56. Gao X, Lin H, Revanna K, Dong Q. 2017. A Bayesian taxonomic classification method for 16S rRNA gene sequences with improved species-level accuracy. BMC Bioinformatics 18:247.

57. Nilsson RH, Larsson KH, Taylor AFS, Bengtsson-Palme J, Jeppesen TS, Schigel D, Kennedy P, Picard K, Glöckner FO, Tedersoo L, Saar I, Kõljalg U, Abarenkov K. 2019. The UNITE database for molecular identification of fungi: handling dark taxa and parallel taxonomic classifications. Nucleic Acids Res 47:D259–d264.

58. Altschul SF, Gish W, Miller W, Myers EW, Lipman DJ. 1990. Basic local alignment search tool. J Mol Biol 215:403–10.

59. Camacho C, Coulouris G, Avagyan V, Ma N, Papadopoulos J, Bealer K, Madden TL. 2009. BLAST+: architecture and applications. BMC Bioinformatics 10:421.

60. Letunic I, Bork P. 2021. Interactive Tree Of Life (iTOL) v5: an online tool for phylogenetic tree display and annotation. Nucleic Acids Res 49:W293–w296.

